# A New High-throughput Screening (HiTS) Method for Phages – Enabling Crude Isolation and Fast Identification of Diverse Phages with Therapeutic Potential

**DOI:** 10.1101/2020.03.27.011080

**Authors:** Nikoline S. Olsen, Niels Bohse Hendriksen, Lars H. Hansen, Witold Kot

**Author notes:** W. K. and L. H. H. initiated and supervised the project equally.

## Abstract

Bacteriophage therapy and application of phages as biocontrol in plant production and food processing, all necessitates acquisition of suitable phages. Depending on purpose, the selection criteria of phage characteristics include lifestyle (lytic/lysogenic), host range, physical stability and absence of unwanted genetic traits such as integrases, antibiotic resistance or bacterial virulence factors. The exclusivity of antibiotic resistant clinical infections and possible development of phage-resistance instigates a need to continually build sizeable phage libraries and also be able to rapidly isolate and characterise novel phages of specified bacterial hosts. Current methods for phage isolation are both laborious and time consuming, suitable only for the isolation of a limited number of phages. Thus, we developed the ***Hi****gh-****T****hroughput* ***S****creening* (HITS) method for phages for fast isolation and identification of potentially hundreds of distinct phages against single hosts. This scalable method enables screening of hundreds of samples, in multiple simultaneous setups with varying parameters increasing the likelihood of isolating multiple distinct phages specific for the given conditions. The efficiency of the method is emphasised by our screening of 200 environmental samples, resulting in the identification of an abundance of unique phage species lytic to *Escherichia coli, Salmonella Enterica, Enterococcus faecalis* and *Pseudomonas aeruginosa*.

## 1. Introduction

The upsurge of antibiotic resistant bacteria is one of the main health concerns of our time [1]. Pathogenic bacterial infections are becoming ever more difficult to treat, and even last resort antibiotics such as the glycopeptide antibiotics vancomycin and teicoplanin are falling short as efficient antimicrobial agents [2]. Bacteria are consecutively acquiring antibiotic resistance and develop multidrug resistance [3], which necessitates the development of alternative antimicrobials or means to increase the efficiency of existing antibiotics. Phage therapy (PT) is the therapeutic use of the viral antagonists of bacteria, the bacteriophages (phages), to treat bacterial infections in humans or animals. Most bacteriophages have narrow host-ranges, limiting their infectivity to specific species or even strains. Consequently, PT does not instigate drastic perturbations of natural microbiota like traditional antibiotic treatments [4]. Though studies have been limited, PT has not been shown to have any adverse side effects [5]. Moreover, PT has shown potential as a last resort treatment of multi-resistant bacterial infections, when traditional antibiotics fall short [6]–[8]. Hence, PT is, especially when applied as a combination therapy together with conventional antimicrobials, foreseen to play an essential role in the multifaceted strategy required to combat the lurking antibiotic crisis [1], [9]. Furthermore, the use of phages for biocontrol in plant production and food processing has displayed a promising potential [10], and could be a sustainable alternative to traditional chemicals facing restrictions due to concerns for public health and the spread of resistance [11].

Yet, a successful biocontrol or PT venture requires phages with different modes of action, and lots of them. Infection-specific phages and prepared phage cocktails are rarely generalisable [12]. Clinical infections can be unparalleled and call for *de novo* isolation or genetic engineering, as was the recent case with a 15-year old patient with cystic fibrosis caused by *Mycobacterium abscessus* [6]. More than 10 000 phages infecting *Mycobacterium smegmatis* were screened in addition to 100 environmental samples, resulting in only three suitable phages, two of them requiring genetic engineering [6]. Indeed, one of the greater hurdles for effective PT, is the availability of suitable phages [12]. Methodologies for isolation of phages have not changed much since phages were discovered more than 100 years ago. The procedures are laborious and time-consuming. In general, phages are isolated by either direct plating or by enrichment and then subsequent purification. Enrichment entails an introduction of a host to a phage-containing sample, which is afterwards removed by centrifugation and filtration when phages have been amplified. Direct plating and purification is typically performed with the soft-agar overlay technique, first described by A. Gratia in 1936 [13]. Improvements to increase throughputs have been proposed, such as tube-free agar overlays [14], and phage activity can now be measured by means more suitable for automation, like colorimetric methods [15]. However, no truly high-throughput isolation method has, to our knowledge been offered. A citizen science approach, like the great effort performed by The Science Education Alliance Phage Hunters Advancing Genomics and Evolutionary Science (SEA-PHAGES) has resulted in the isolation of thousands of phages against *M. smegmatis* [16]. But this type of approach requires both substantial funding and facilities.

In order to establish and expand libraries of phages relevant for PT and biocontrol, affordable, fast and efficient screening methods are needed to enable rapid isolation and identification of candidate phages. Large libraries of phages infecting the same single host, also enables important phage-host interaction studies, expanding our understanding of phage taxonomy and ecology. Here we present the ***Hi****gh-****T****hroughput* ***S****creening* (HiTS) method for phages, which enables a single person to go from a high number of samples to a plethora of identified phages within weeks. The simplicity of the method enables >500 samples to be handled simultaneously. The HiTS method selects for predominantly lytic and easily culturable phages. The resolution is a single or a few phages from each sample processed. The integrated sequencing of the identified phages allows for an early assessment of genomes enabling the selection of candidates which do not possess any unwanted genetic-traits and are thus suitable for further characterisation and potential application as PT or biocontrol agents.

## 2. Materials and Methods

The method presented is host-system independent and can thus be applied for screening of environmental samples for phages virulent to any culturable fast-growing aerobic or facultative anaerobic bacteria by adjusting host media, incubation temperature and time. The protocol enables a simple and fast (4 consecutive days), yet crude purification of single or a low number of distinctive phages from a small sample volume (0.5-1.5 ml). The method allows for a high number of samples to be handled, with simultaneous investigation of diverse sample matrices or parallel screenings of the same sample-set with varying parameters e.g. host, pH, media, amendments and incubation conditions (Figure 1). This increases the likelihood of sequestering multiple distinctive phages from each sample. The method is suitable for both direct plaque sequencing (DPS) [17] and standard phage DNA extraction from lysate. The screening procedure entails four steps: 1. *Phage amplification, 2. Liquid purification*, 3. *Spot-test* and 4. *Phage collection and DPS or optional: plating of dilution series*.

**Figure 1.**
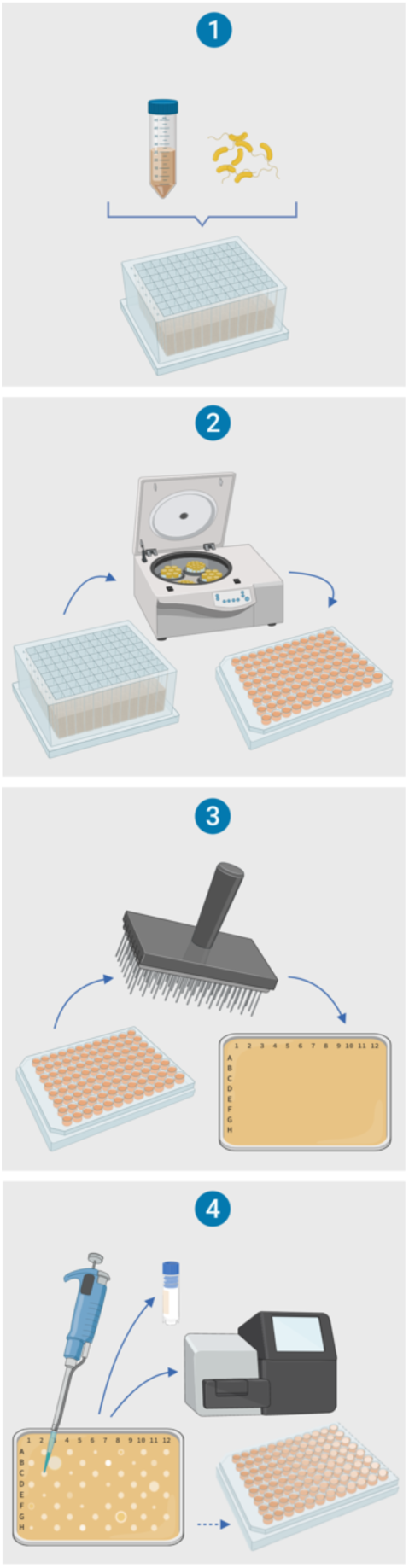
The four steps in the HiTS method. Illustration created with BioRender.

### 2.1. Protocol

#### High-throughput Screening (HiTS) Method for Phages

The method, which is scalable for robotics, requires a centrifuge suitable for 96-well plates and a 96-pin replicator or a one µl multichannel pipette. Multichannel pipettes or a pipetting robot may ease many of the steps involved. Dry samples should be suspended prior to processing and preferably centrifuged and filtered. The maximum number of samples per plate is 94. The sample volume can be adjusted as desired and as applicable to available well plates. By increasing the volume of raw sample input, the number of phages per incubation is also increased and thus the chance of isolating these. Initial sample volume only affects step *1. Phage amplification*. Volumes and concentrations suggested in step 0. and step 1. are suitable for screening 1.5 ml raw sample in deep-well plates with a working volume (wV) of 2.2 ml (e.g. 732-0612, VWR, Radnor PA US). All incubations should be performed under optimal host conditions (media and temperature) and hence adjusted as required. In step 4 there is the option to either collect the phages and sequence them by DPS or to do titers and aim for single plaques, and then do DPS or sequence phage-amplifications.

#### 0. Preparation

a. Prepare all media, solutions and agar-plates.
b. Inoculate host cells in 2 × 10 ml liquid media and incubate overnight
c. (ON).

##### 1. Phage amplification (day 1)

a. Distribute a maximum of 94 samples, in suitable volumes (1.5 ml) in a deep-well plate (#1) with pierceable sealing tape (e.g. Z722529-50EA, Excel Scientific, Victorville CA US). Sterilised water (1.5 ml) is added as negative amplification-controls to wells D6 and E6. To each of the 96 wells add:

90 µl CaCl_2_ (0.25 M) and MgCl_2_. (0.25 M), final conc. 10 mM
110 µl ON host culture, final conc. 5% V/V
500 µl Host media (conc. × 4.4), final conc. × 1 During addition of media, carefully pipette up and down a few times to mix. Close the well-plate and incubate ON on a shaker (200 rpm).
b. Inoculate ON host culture in 10 ml liquid media for next day.

##### 2. Liquid purification (day 2)

a. Filter to remove host bacteria by transferring 200 µl (punch through pierceable tape) from each well to a 96-well filter Plate (0.45 µm) (e.g. MSHAS4510, Merck Millipore, Burlington MA US), pipette up and down a few times before extracting. Centrifuge filter plate on top of a new well-plate (#2, wV 200 ul e.g. 269787, Nunc, Roskilde, DK) at 900 x *g* for 2 minutes. Then add pierceable sealing tape to well-plate #2. Discard the filter plate.
b. Prepare a third well-plate (#3, wV 200 µl) with pierceable sealing tape and add:

180 µl host media (conc. x 1)
10 µl of host culture, final conc. 5% V/V
10 µl 0.2 M CaCl_2_ and 0.2 M MgCl_2,_ final conc. 10 mM Use the 96-pin replicator to transfer ∼1 µl of each lysate (punch through pierceable tape) in well-plate #2 to each well in well-plate #3. Close well-plate #3 and incubate ON on a shaker (200 rpm). If processing more than one set of samples, clean the 96-pin replicator by ethanol and flame three times in between and make sure to cool it down before re-use.
c. Inoculate ON host culture in 10 ml liquid media for next day.

##### 3. Spot-test (day 3)

a. Filter to remove host bacteria as described in step 2a.
b. Prepare two large (Ø 14 cm e.g. 82.1184.500, Sarstedt, Nürnbrecht DE or at least 12 × 8 cm e.g. 242811, Nunc) soft-agar overlay plates (A and B) of 0.5% agarose amended with: CaCl_2_ and MgCl_2_ (final conc. 10 mM) Host culture (final conc. 2.5 - 5 %)
c. While the plates solidify remove every second row of pipette-tips in a box of 200 µl pipette-tips to facilitate the transfer of lysate from every second well in a chequered pattern into two new microtiter plates with pierceable sealing tape, number #4 (A) and #5 (B) (Figure S1).
d. Use the 96-pin replicator to carefully transfer ∼l µl of lysates from well-plate A (#5) to the softagar overlay plate A. Make sure to specify direction on the plate. The chequered pattern ensures a safe distance between spotted samples, a negative amplification-control (D6 or E6) on each plate and sterilisation-controls (every second tip) (Figure 1). Clean the 96-pin replicator by ethanol and flame and repeat the procedure with well-plate B (#5) and soft-agar overlay plate B. Incubate soft-agar overlay plates upside down ON. Seal well-plates A (#4) and B (#5) and store at 4°C.
e. **Optional**: Inoculate ON host culture in 10 ml liquid media for titre estimation next day.

##### 4. Phage collection and DPS, or optional: plate dilution series (day 4)

a. The centre of clearing zones (agar-plate A and B) is collected for DPS. Additional clearing zone is dissolved in 100 µl SM-buffer [18], filtered (0.22 - 0.45 µm) and stored for future purification and characterization. If clearing zones are too small for double collection make amplifications of the phage-SM solutions (inoculate host bacteria in 10 ml media, after ∼1 h add lysate, next day centrifuge and filtrate) and extract DNA for sequencing from these.
b. Optional: titre the lysates by transferring phage solutions from positive (plaque-forming) wells to new wells in column 1 of as many new well-plates (#6+, wV 200 µl) as required. Eight-fold dilutions series are made within the well-plates by adding 180 µl SM-buffer to all wells in column 2-9 and then transferring 20 µl of the solution in column 1 to column 2 pipetting up and down to mix and repeating the procedure for the remaining columns. Spot (∼1 µl) the dilution series on soft-agar overlay plates with a 96-pin replicator or multichannel pipette and incubate ON. Next day: Count plaques or clearing zones for approximate titre. Do DPS of single plaques if present and also collect clearing zone for phage storage. If single plaques are not present plate lysate dilution giving rise to 10-50 plaques on a full plate by the soft-agar overlay method (add the lysate to 4 ml 0.5% agarose with 10 mM CaCl_2_ and MgCl_2_ and 2.5-5% (V/V) ON host culture, pour on standard petri dish with agar). Next day, pick diverting plaque morphologies for DPS or phage-amplification, lysate hereof can be used for DNA extraction and phage storage.

### 2.2. Phage screenings

Five screenings were performed as described in *2.1 Protocol*, using *Escherichia coli, Salmonella enterica, Enterococcus faecalis* or *Pseudomonas aeruginosa* as hosts (Table 1). For the *E. coli, E. faecalis* and *S. enterica* screenings 188 distinct wastewater samples divided into two sets of 94 samples were used, for the *P. aeruginosa* screening 82 wastewater samples were used together with eight soil samples and four organic waste samples (Table S1). The *S. enterica* screenings were performed with both a small sample volume (SV) of 0.5 ml and a large sample volume (LV) of 1.5 ml, while the *E. coli* and *E. faecalis* screenings were only performed with 0.5 ml (SV) and the *P. aeruginosa* screening only with 1.5 ml (LV). The SV screenings (*E. coli, E. faecalis* and *S. enterica)* followed the protocol, with the exception that instead of DPS, lysates from step 3. corresponding to positive wells (those instigating clearing zones) were used for DNA extraction and sequencing, while phages were stored by collecting top-agar of clearing zones or plaques. In step 4 of the LV screenings (*S. enterica* and *P. aeruginosa*) lysates from positive wells were tittered in the 96-well format and the most diluted lysates instigating single plaques or clearing zones were used for making 10 ml amplification lysates for DNA extraction, library preparation, sequencing and phage storage. All incubations were performed at 37°C.

**Table 1.** Number of samples screened, clearing zones detected and phages identified in the five HiTS screenings, using *S. enterica, E. coli, E. faecalis* or *P. aeruginosa* as host. *Escherichia* phage data from Olsen *et al*., (2020) [39].

| Host | Sample (ml) | Samples (n) | Clearing zones (n) | Sequenced lysates<sup>1</sup> (n) | Phages (n) | Unique species<sup>2</sup> (n) | Novel species<sup>3</sup> (n) |
| --- | --- | --- | --- | --- | --- | --- | --- |
| *S. enterica* | 0.5 SV | 188 | 51 | 42 | 47 | 33 | 28 |
| *S. enterica* | 1.5 LV | 188 | 74 | 60 | 76 | 45 (26<sup>4</sup>) | 38 (24<sup>4</sup>) |
| *E. coli* | 0.5 SV | 188 | 153 | 94 | 136 | 104 | 91 |
| *E. faecalis* | 0.5 SV | 188 | 5 | 4 | 4 | 4 | 3 |
| *P. aeruginosa* | 1.5 LV | 94<sup>5</sup> | 48 | 38 | 46 | 22 | 8 |

<sup>1</sup>Includes all lysates for which sequencing yielded reads assembling to contigs with a coverage $> \times 20$ . <sup>2</sup>Phages with $\leq 95$ % nucleotide similarity to the other phages in this dataset. <sup>3</sup>Phages with $\leq 95$ % similarity to the other phages in this dataset and those deposited in the NCBI database. <sup>4</sup>Excluding the phages with $>95$ % nucleotide similarity to phages in the *S. enterica* SV screening. <sup>5</sup>82 wastewater samples, 8 soil samples and 4 organic waste samples.

#### 2.2.1. Bacteria and growth media

The host bacteria used for phage screenings were *E. coli* (K-12, MG1655), *S. enterica subsp. enterica* serovar Enteritidis PT1, the vancomycin-resistant *E. faecalis (*strain ATCC 700802 / V583) and the chloramphenicol-resistant *P. aeruginosa* (PAO1). The media applied was LB (Alpha Biosciences, Baltimore MD US).

#### 2.2.2. Samples

The 188 inlet wastewater samples (40-50 ml) were collected in time-series of 2-4 days during July and August 2017, from 48 Danish wastewater treatment facilities geographically distributed in both rural and urban areas on Zealand, Funen and in Jutland. Upon receipt, the samples were centrifuged (9000 x g, 4 °C, 10 min), the supernatant filtered (0.45 µm) and then stored in aliquots at -20°C. The organic waste samples were collected from four different Danish facilities in February, May and November 2017. The 12 soil samples (∼5 g) were collected in Roskilde municipality, Denmark, in March 2019. Prior to screening the soil was suspended in 5 ml LB and slowly and continuously inverted for 1h at room temperature, then the samples were centrifuged (9000 x g, 5 min) and the supernatant filtrated (0.45 µm). Refer to Table S1 for a list of all samples and facilities.

#### 2.2.3. DNA extraction, library preparation and sequencing

Phage DNA extractions were performed by an initial DNase treatment, 1 U of DNase 1 (New England Biolabs, Ipswich, MA US) per ∼100 µl lysate (37°C, 30 min, inactivated by 10 µl 50 mM EDTA), followed by addition of 3 U Proteinase K (A & A Biotechnology, Gdynia, Poland) and 10% (v/v) SDS solution (55°C, 30 min, inactivated by 70°C, 10 min). The extracted DNA was then purified in the well-plate format using the ZR-96 Clean and Concentrator kit (Zymo research, Irvine, CA US), following manufacturer’s instructions and eluting in 6 µl of the supplied elution buffer. Sequencing libraries were built according to manufacturer’s instructions with minor modifications as described in Kot *et al*., (2014) [17] using the Nextera® XT DNA kit (Illumina, San Diego, CA USA), the libraries were sequenced as paired-end reads on Illumina NextSeq platform with the Mid Output Kit v2 (300 cycles).

#### 2.2.4. Assembly, annotation, identification and phylogenomic analysis

The obtained reads were trimmed and assembled in CLC Genomics Workbench 10.1.1. (CLC BIO, DK), overlapping reads were merged with the following settings: mismatch cost: 2, minimum score: 15, gap cost: 3 and maximum unaligned end mismatches: 0, and then assembled *de novo*. Additional assemblies were constructed using SPAdes 3.12.0 [19]. Gene prediction and annotation was performed using a customized RASTtk version 2.0 [20] workflow with GeneMark [21], with manual curation and verification using BLASTP [22], HHpred [23] and Pfam version 32.0 [24], or *de novo* annotated using VIGA version 0.11.0 [25] based on DIAMOND searches (RefSeq Viral protein database) and HMMer searches (pVOG HMM database). NT similarity was determined as percentage query cover multiplied by percentage NT identity. Novel phages were categorised according to ICTV taxonomy. The criterion of 95% DNA sequence similarity for demarcation of species was applied to identify novel species representatives and to determine species uniqueness within the dataset. All unique phage genomes were deposited in GenBank (Table 1). All genomes were assessed for antibiotic resistance genes (ARGs) and bacterial virulence genes using ResFinder 3.1 [26], [27] and VirulenceFinder 2.0 [28]. NT and amino acid (AA) similarities were calculated using tools recommended by the ICTV [29], i.e. BLAST [22] for identification of closest relatives (BLASTn when possible, discontinuous megaBLAST (word size 16) for larger genomes) and Gegenees version 2.2.1 [30] for assessing phylogenetic distances of multiple genomes, for both NTs (BLASTn algorithm) and AAs (tBLASTx algorithm) a fragment size of 200 bp and step size 100 bp was applied. Evolutionary analyses for phylogenetic trees were conducted in MEGA7 version 2.1 (default settings) [31]. These were based on the large terminase subunit *terL*, a gene commonly applied for phylogenetic analysis [32], [33] and on the DNA encapsidation gene *gpA* for the <13kb alleged *Podoviridae*. The NT sequences were aligned by MUSCLE [34] and the evolutionary history inferred by the Maximum Likelihood method based on the Tamura-Nei model [35]. The tree with the highest log likelihood is shown manually curated by adding color-codes and identifiers in Inkscape version 0.92.2. The R package iNEXT [36], [37] in R studio version 1.1.456 [38] was used for rarefaction analyses, species diversity (q = 0, datatype: incidence_raw), extrapolation hereof (estimadeD) and estimation of sample coverage. Additional graphs were prepared in Excel version 16.31.

## 3. Results

### 3.1. Screening efficiency and resolution

Across all five screening between 3% (n = 5 of 188) and 81% (n = 153 of 188) of samples yielded clearing zones plausibly due to lysis by phages, the majority of these also gave rise to the identification of phages (Table 1). However, in some cases the DNA extraction was unsatisfactory or the sequencing failed. Between 61% (*E. coli* screening n = 94 of 153) and 82% (*S. enterica* SV screening n = 42 of 51) of clearing zones were successfully sequenced i.e. yielded reads assembling to phage contigs with an average coverage > x 20 (Table 1). Regardless of host, a single phage was identified from the vast majority of sequenced samples (64-100% per screening), although in some instances two (0-29% per screening), three (0-6% per screening) or four (0-1% per screening) phages were identified from a single sample (Figure 2a). The *Escherichia* phages were the most numerous (136 phages from 94 wells), they were more frequently (34 samples) isolated as more than one phage per sample and the only ones to be four phages in a sample [39] (Figure 2a, Table 1). The number of phages per sample did not differ considerably between phages of *S. enterica* (123 phages form 102 wells) and *P. aeruginosa* (43 phages from 38 wastewater wells), while only four phages lytic to *E. faecalis* were identified in four separate samples (Figure 2a). *Escherichia* phages were identified in samples from 43 different facilities out of the 48, *Enterococcus* phages in samples from 4 facilities and *Salmonella* phages in samples from 22 of the 48 facilities included in these screenings. In the *P. aeruginosa* phage screening, phages were identified in wastewater samples from 95% of the 21 facilities included, (20 out of 21). Furthermore, *P. aeruginosa* phages were identified in three of the four organic waste samples, but in none of the 8 soil samples (Figure 2c).

**Figure 2.**
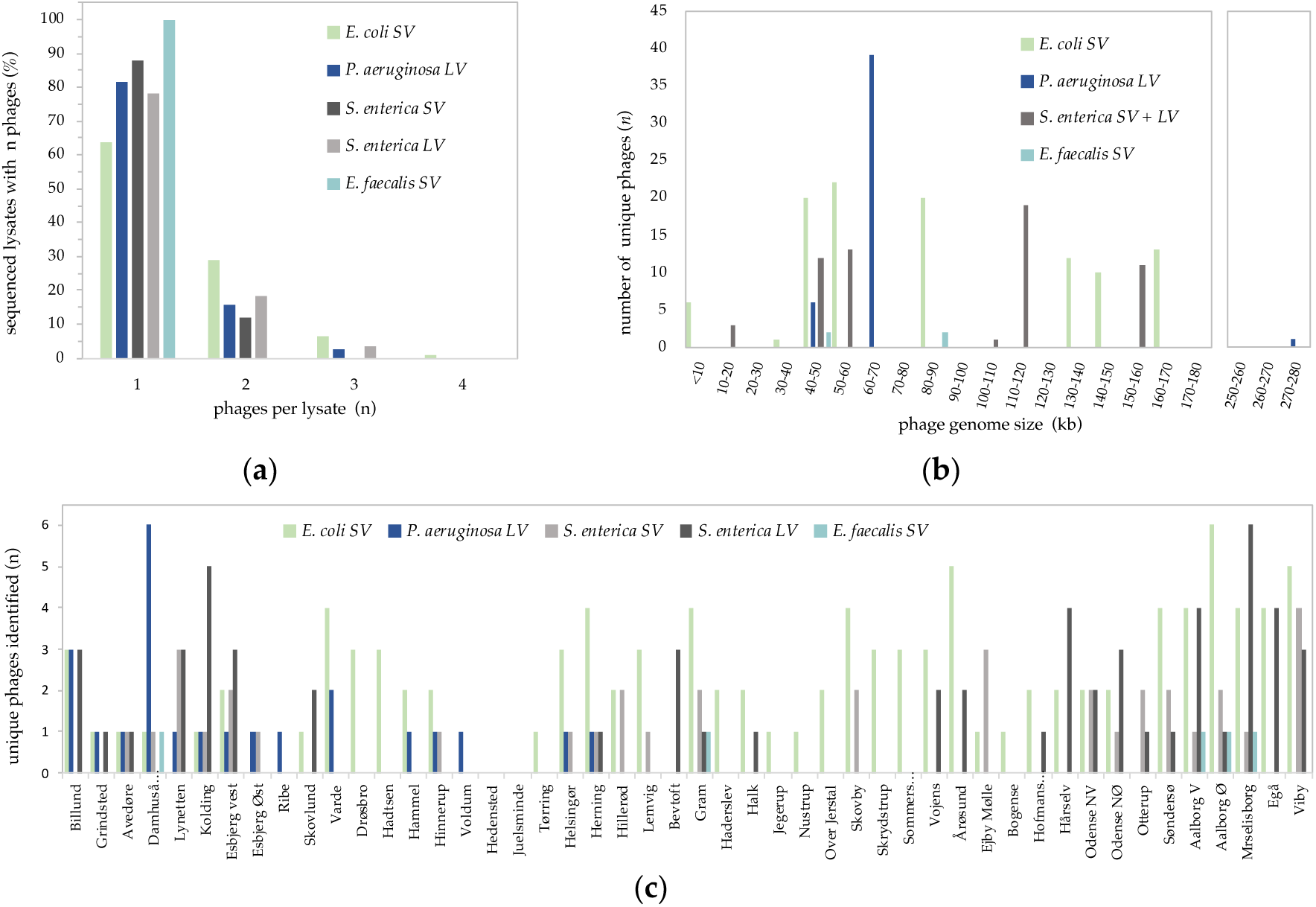
(**a**) Number of phages per lysate (n) (x-axis) as occurring in percentage of all sequenced lysates (y-axis), presented according to individual screenings. (**b**). Average genome size (kb) of Gegenees based clusters, of unique phage species (<95% nucleotide similarity to other phages in the dataset) organized by host, from all screenings. (**c**) Distribution of all 309 phages identified organised per facility, only the first 21 facilities were included in the *Pseudomonas aeruginosa* screening.

Of the 136 *Escherichia* phages, the majority (76%) represent unique species [39]. The many *Salmonella* and *P. aeruginosa* phages are more homogeneous. The two *S. enterica* phage screenings identified 123 phages. Out of 47 SV phages 14 were shown to have >95% NT similarity with other SV phages, while 31 of the 76 LV phages were shown to have >95% NT similarity with other LV phages and an additional 19 of the LV phages had >95% NT similarity with SV phages. Hence, a total of 59 (48%) *Salmonella* phages of distinct species are identified. Similarly, of the 46 *P. aeruginosa* phages 22 (48%) are unique, while all four *E. faecalis* phages represent distinctive species (Table 1).

### 3.2. Novelty and diversity of HiTS-phages identifed compared to NCBI

The phages identified cover an impressive wide range of genome sizes (Figure 2b, 4), GC contents and predicted morphologies, representing five different families; the *Ackermannviridae, Myoviridae, Podoviridae* and *Siphoviridae* of the order *Caudovirales* and also the non-tailed *Microviridae* (Table 2) [39]. The *Escherichia* phages are indeed remarkably numerous and diverse and are consequently described separately in Olsen *et. al*. (2020) [39]. In summary, disregarding the jumbo phage Pseudomonas phage fnug (278.9 kb), the *Escherichia* phages cover the largest size range (5.3-170.8 kb) and have an impressive GC content span (35.3-60.0%) [39]. Members of the new family *Ackermannviridae*, were only detected among the *Salmonella* phages, just as members of the *Microviridae*, a family of small single stranded DNA phages, were only observed among the *Escherichia* phages [39]. The *Salmonella* and *P. aeruginosa* phages covered similar GC content spans of 36.9-56.5% and 39.0-59.5%, respectively. The *Salmonella* phage genomes vary in sizes from 11.6 kb (Salmonella phage astrithr) to 159.1 kb (Salmonella phage maane), while the non-jumbo *P. aeruginosa* phages are more uniform having genome sizes of 44.9 kb (Pseudomonas phage clash) to 66.5 kb (Pseudomonas phage shane). The *Enterococcus* phages have genomes of 39.7-85.7 kb, Enterococcus phage heks and nattely, respectively (Figure 2b, Tables S2-S4).

**Table 2.** Predicted morphology of all the phages identified in screenings for *Escherichia, Salmonella, P. aeruginosa* and *Enterococcus* phages, based on taxonomy of closest relative. *Escherichia* phage data from Olsen *et al*., (2020) [39].

| Phage taxonomy | Isolation hosts: | *E. coli* | *S. enterica* | *P. aeruginosa* | *E. faecalis* |
| --- | --- | --- | --- | --- | --- |
| *Caudovirales; Ackermannviridae* |  | - | 21 | - | - |
| *Caudovirales; Myoviridae* |  | 79 | 18 | 39 | - |
| *Caudovirales; Siphoviridae* |  | 34 | 76 | 1 | 4 |
| *Caudovirales; Podoviridae* |  | 10 | 8 | 6 | - |
| *Microviridae; Bullavirinae* |  | 8 | - | - | - |
| *Unclassified bacterial viruses* |  | 5 | - | - | - |

An impressive number (n = 154) of novel phage species have so far been identified with the HiTS method. No less than 67% (n = 91) of the *Escherichia* phages [39], 42% (n = 52) of the *Salmonella* phages, 22% (n = 8) of the *P. aeruginosa* phages and three out of four *Enterococcus* phages represent novel phage species (Table 1, Figure 3). Whereas most of the *Escherichia* (69%) and all of the *P. aeruginosa* phage species representatives have a high NT similarity (>89%) with their closest relatives, a larger proportion of the *Salmonella* phages differ more from their closest relatives as only 54% (n = 42) of the unique *Salmonella* phages species have >89% NT similarity to their closest relative (Figure 3). Two of the *Escherichia* phages and three *Salmonella* phages share <50% NT similarity with published phages. Likewise, two of the *Enterococcus* phages (Figure S3, Table S3) and Salmonella phage Akira (63% NT similarity) are only distantly related to any published phage (62-65% NT similarity) (Figure 3). The 189 unique phages (<95% NT similarity with other phages in the dataset) and their GenBank accession numbers are listed in supplementary materials (Table S2-S5).

**Figure 3.**
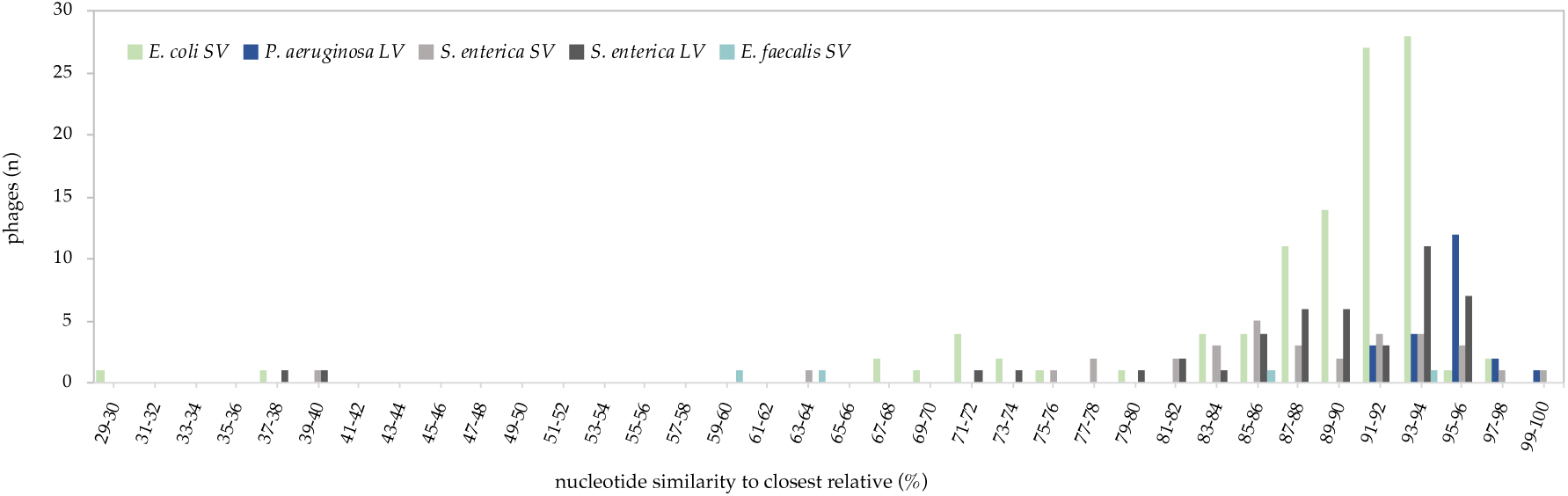
Distribution of nucleotide similarity (%) to closest relative (Blast) of all phages from all five screenings.

No virulence factors or ARGs were detected in any of the 189 unique phages. Furthermore, no integrases were identified and though putative recombinases do occur it is uncertain if they are involved in DNA repair or integration.

### 3.3 P. aeruginosa and Enterococcus phages from Danish wastewater

The *P. aeruginosa* phages group into two distinct clusters and two singletons (Figure S2). The vast majority (n = 38) of the *P. aeruginosa* phages are closely related (90.9-98.5% NT similarity) to phages of the genus *Pbunavirus* of the family *Myoviridae*, with genome sizes of 60.7-66.8 kb (89-95 CDSs, 54.8-55.7% GC, no tRNAs,). A smaller group of six *P. aeruginosa* phages (44.9-45.3 kb, 68-69 CDSs, 52.1-52.5% GC, 3-4 tRNAs) are closely related (94.1-96.3% NT similarity) to phages of the genus *Bruynoghevirus* of the family *Podoviridae*. The jumbophage fnug is closely related (93% NT similarity) to phages of the genus *Phikvirus*, family *Myoviridae*. While the last Pseudomonas phage Iggy (60.7 kb, 90 CDSs, 56.5% GC, no tRNAs) is closely related (94.6%) to the unclassified *Siphoviridae* Pseudomonas phage PBPA162 (MK816297), none of them share >8% NT similarity with any other published phages. The *Enterococcus* phages are all predicted to have *Siphoviridae* morphology, but divide into two distinct clusters with NT inter-Gegenees scores of 0 (Figure S3). Phages heks and Nonaheksakonda (39.7-41.9 kb, 64-74 CDSs, 34.6-35.0% GC, no tRNAs) are related to efquatroviruses, but with only 59% NT similarity. The other two (85.3-85.7 kb, 131-134 CDSs, 30.2-30.3% GC, 1 tRNA) are more closely related (87-96% NT similarity) to unclassified *Siphoviridae* (Figure 2b, Tables S2-S5).

### 3.4 Salmonella phages from Danish wastewater

Based on NT similarity with closest relatives, 59 distinctive species of *Salmonella* phages were identified, of which 52 represent novel species. Estimations based on both the SV and LV screenings predicts species richness of easily culturable phages lytic to *S. enterica subsp. enterica* serovar Enteritidis PT1 in Danish wastewater to be nearby 80 (Figure S4), while Shannon diversity estimates 68 and 61 and Simpson diversity 51 and 38, for the SV and LV screenings, respectively (Figure S4). The estimates are however expected to be subject to large prediction bias due to the relatively small reference sample size, and a 95% confidence interval suggests a range for all diversity indices of 26-173. Sample completeness is estimated to be achieved at ∼1300 samples for a SV screening and at ∼800 samples for a LV screening (Figure S4). The HiTS *Salmonella* phages belong to at least four different families, *Ackermannviridae, Myoviridae, Podoviridae* and *Siphoviridae*, covering a wide range of genome sizes and GC contents (Table S5). They group into five clusters and three singletons (inter-Gegenees scores = 0) corresponding to their proposed taxonomy, excluding phage Akira (Figure 4). In spite of a NT similarity of 63% and a Gegenees score of 38-39 with its closest relative the unclassified *roufvirus* Salmonella virus KFS_SE2 (MK112901) Akira does not group with neither KFS_SE2 or the type species of *roufvirus* Aeromonas phage pIS4-A (NC_042037) in the phylogenetic tree, resulting in a peculiar pattern in the Gegenees analysis (Figure 4). Furthermore, Akira shares limited NT similarity (39%) with pIS4-A and a Gegenees score of only 3-4 (Figure 4, Table 2).

**Figure 4.**
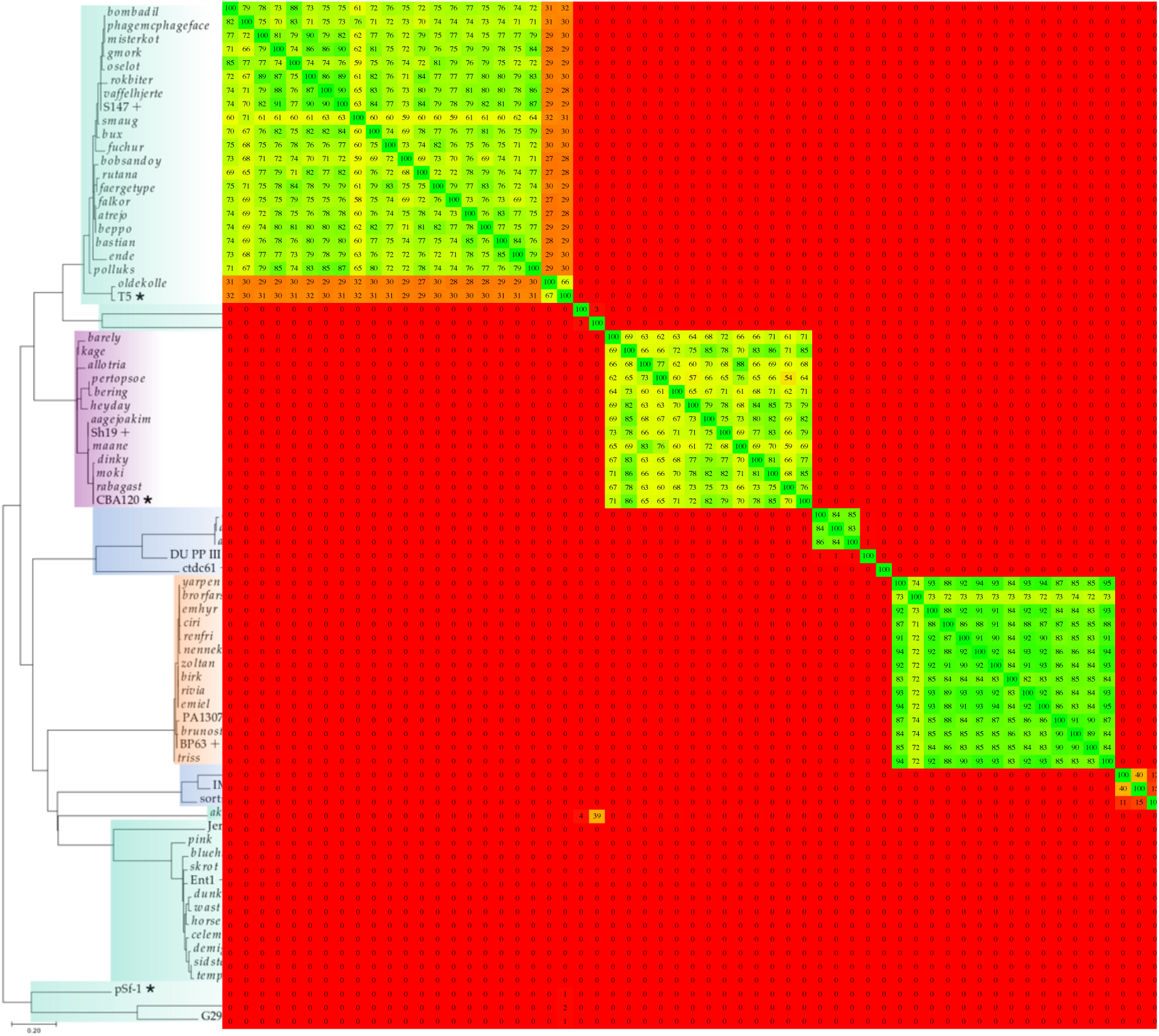
Phylogenetic tree (Maximum log Likelihood: -4176.67, based on large terminase subunit or for the <13 kb *Podoviridae* the DNA encapsidation protein, scalebar: substitutions per site) and phylogenomic nucleotide distances (Gegenees, BLASTn: fragment size: 200, step size: 100, threshold: 0%). Predicted morphology is indicated by colorbars, ▪ *Myoviridae*, ▪ *Siphoviridae*, ▪ *Podoviridae*, ▪ *Ackermannviridae*, novel phages from this study are in italics, while close relatives are denoted by + and type species by *

All nine novel phages (143-159 kb, 187-209 CDSs, 44-45% GC, 3-4 tRNAs) of the *Ackermannviridae* are related (78-99% NT similarity) to phages of the genus *Kuttervirus*, subfamily *Cvivirinae*, defined by an average genome size of 158.1 kb, with and average GC content of 44.5% and averagely coding for 201 proteins and 4.3 tRNAs. These nine phages have an average NT intra-Gegenees score of just 60, but are more similar to one another and also to the type species of *Kuttervirus*, Escherichia virus CBA120 (JN593240) when comparing AAs (Gegenees score = 88-97).

Most of the *Salmonella* phages are predicted to belong to *Siphoviridae* (n = 76, 32 species, 29 novel species). Twenty of the unique *Siphoviridae* phages (105-115 kb, 151-171 CDSs, 39.3-40.1 GC, 23-31 tRNAs) are related (71-95% NT similarity) to phages of the genus *Tequintavirus*, the T5 like phages. However, whereas phage oldekolle has low NT Gegenees scores with the rest (n = 27-32) and is closely related to T5 (93% NT similarity), the remainder appear more distantly related to T5. Even though, these phages cluster together, they have relatively low NT inter-Gegenees scores (n = 59-90) and they also differ from published phage genomes by 6-29% NT similarity (Figure 4, S5, Table S5). An additional ten unique *Siphoviridae* phages (41-44 kb, 56-70 CDSs, 50% GC, 0-1 tRNA) were found to be closely related to jerseyviruses, subfamily *Guernseyvirinae*. This genus is defined by genome sizes of 40-44 kb, comparable morphology and a shared DNA identity of ≥60% and >68% protein content [40]. The wastewater jerseyviruses-like phages are a heterogeneous group with varying NT intra-Gegenees scores of 38-91, yet the AA intra-Gegenees scores are all >69. However, the Gegenees NT scores between the novel phages and the type species Salmonella phage Jersey (NC_021777) are all <33 though the AA scores are 65-71. The novel jerseyviruses-like phages are relatively abundant in the Danish wastewater samples and most of them were identified in several samples from different treatment facilities. Phages with >95% NT similarity with phage wast were found eight times in samples from five distinct facilities and phages with >95% NT similarity with phage demigod as many as 12 times, in samples from six different facilities (Table S5).

The last *Siphoviridae* within the *Salmonella* phages is the novel phage slyngel, related (92% NT similarity) to Escherichia phage vB_EcoS_G29-2 (MK373798) an unclassified *Hanrivervirus*, subfamily *Tunavirinae*. Furthermore, slyngel has 88% NT similarity with the type species Shigella phage pSf-1 (NC_021331), with which a more distant relationship is also suggested by the phylogenetic analysis (Figure 4, S6).

Twelve of the *Salmonella* phages are based on NT similarity of the *Myoviridae* and though their closest relatives are all unclassified *Myoviridae*, they constitute the most homogeneous group. They have comparable genomes (52-53 kb, 67-73 CDSs, 45.7-46% GC, no tRNAs) and cluster together in both the Gegenees analyses and in the phylogenetic tree. Though phage brorfarstad has slightly lower NT Gegenees scores (n = 72-74), than the rest, only a minor difference can be observed in the AA Gegenees analysis (Figure 4, S5, Table 3).

The eight *Podoviridae* divide into a cluster of three phage species representatives (astrid, assan and astrithr) with comparable (83-84 NT Gegenees score) small low GC genomes (11.6-11.7 kb, 15 CDSs, 39.7-39.8% GC, no tRNASs) and the singleton lumpael. Phages with >95% NT similarity with phage Lumpael were observed in five samples from five distinct facilitates. Lumpael (41.1 kb, 58 CDSs, 59.9% GC, no tRNAs) has the highest GC content observed and shares only 76% NT similarity with its closest relative Enterobacteria phage IME_EC2 (KF591601). Astrid, assan and astrithr all share <40% NT similarity with their closest relative Pectobacterium phage DU_PP_III (MF979562), and though they share NT Gegenees scores of only 0-1 and AA of 45, they do form a monophyletic clade (Figure 4, S5).

## 4. Discussion

When it comes to phage isolation three aspects are key; titre, sterility and purity. These features are a prerequisite for any phage work, regardless of aim. However, purity and sterility are not the main focus in a screening such as the HiTS method. This method does not intend to provide a final PT or biocontrol product, but instead offers crude isolation of highly diverse phages in sufficient titres. This is to provide a fast and crude acquisition of numerous and diverse phages and thereby a basis for further phage isolation and establishment or expansion of phage libraries. If a phage of interest is in a mixed lysate, sequencing enables primer design for PCR verification when isolating the individually plaquing phages.

The method is unique in its ability to identify numerous assorted phages while also facilitating crude isolation in a very short time span. This makes investigations of phage diversity easy and possible. The capacity of the HiTS method to uncover diverse phages is clearly illustrated by the impressive findings when screening for phages of PT relevance in wastewater, especially those infecting *E. coli* and *S. enterica*., as presented in Olsen *et al*. (2020) [39] and in this study, respectively. Even for less abundant phages diversity and novelty was uncovered, the *P. aeruginosa* phages represent three distinct families, and eight are novel phage species representatives, while all the *Enterococcus* phages are of the *Siphoviridae* two of them have limited NT similarity (<60%) with published phages. The reported five screenings yielded no less than 331 potential hits in the form of clearing zones resulting in the identification of 154 novel phage species. Furthermore, none of these phages code for known virulence factors or ARGs and none appear to be lysogenic, making them all potential candidates for PT and biocontrol applications.

Unlike metagenomic sequencing approaches, this method provides actual phages with a direct link to the pathogenic host or any host in interest. The HiTS method does not reveal the diversity of individual samples; hence many phages remain undetected, especially those which are not easily grown under laboratory conditions. The HiTS method is a competition-based method and clearly selects for lytic phages with traits preferable in PT and biocontrol applications i.e. a high burst size and a short latency period. Consequently, the HiTS method enables the capture of the most prevalent phage(s) of the day in any sample. Thus, when screening numerous distinct samples, it provides an estimate of species richness of this type of phages in the given sample matrix. Accordingly, the species richness of easily culturable phages, presumably with high burst-sizes and short latency times, lytic to the specific strains of *Escherichia* and *Salmonella* phages in Danish wastewater was estimated to be at least in the range of 160-420 and 49-173, respectively (Figure S4) [39]. This is likely an underestimation considering the relatively small sample sizes and the inherent bias in the method to only isolate a single or a few phages per sample combined with the many plaque-forming lysates for which DNA-extraction or sequencing was unsuccessful. If the aim of screening is to isolate phages with PT or biocontrol potential or phages that are easy to study under laboratory-conditions, the targeting of lytic and highly reproductive phages is indeed an advantage. However, if the aim is to disclose true diversity or detect more difficult to culture specimen, other methods such as plaquing without amplification may be superior. Metagenomic sequencing approaches are constantly refined and now offers high detection levels of phageomes [41], but phages of interest detected may be near impossible to isolate *in vitro*. The key advantage of the HiTS method is indeed that it offers both identification through sequencing and also provides physical isolates of all phages targeting the specific host-species used as bait. Consequently, interesting discoveries such as rare and novel phages or the presence of remarkable genes with unexpected or desired functions can be investigated following a final isolation. It should however be noted that it is not recommended to sequence the lysate giving rise to plaques or clearing zones, while harvesting phages from the plaques, as was done by the authors in the SV screenings. This approach may result in sequencing of phages in lysate not present in the harvested soft-agar.

A 96-well setup carries a risk of cross-contamination, however the use of pierceable sealing-tape as recommended in the HiTS method, reduces this risk. The presence of a negative amplification control in each spot-test (agar-plates A and B) provides an indication of potential cross-contamination. No plaquing was observed in negative amplification controls in any of the screenings. Furthermore, the chequered-pattern with empty wells between all lysates used when performing the spot-tests ensures that in the case of improper sterilisation, still no phages will be transferred to other wells in use during spotting with a 96-pin replicator, as opposite patterns are present in well-plate A and B. Finally, if the sterilisation of the 96-pin replicator is insufficient any contaminating phages will plaque in between purposely spotted phages. This was not observed in any of the screenings.

A high number of distinct samples, as required by the HiTS method, may be cumbersome to collect and prepare, but once they are collected, they can be aliquoted, stored (−20°C) and used for numerous screenings of different target bacteria, as only very small sample volumes (0.5-1.5 ml) are required. The small sample-volumes also permits for samples to be collected by sending out collection-kits and having them returned by mail or carrier, provided that the applicable law allows it. Any sample matrix with high quantities of the target host is applicable. Furthermore, the suitability of time-series of wastewater-samples eases the sample collection and makes the screening method more feasible as it limits the number of distinct sampling sites. In this study, no *Enterococcus*, only two *P. aeruginosa* (9%), five *Salmonella* (8.5%) and nine *Escherichia* phage species (8.6%) [39] were detected more than once in samples from the same facility in any distinct screening. Wastewater treatment plants receive inlet wastewater in a constant yet changing flow thus the presence of diverse phages can be expected to fluctuate and be interchangeable.

Sequencing is continuously getting cheaper [42], and even though this is the major expenditure of the HiTS method, it is economically feasible. Spending weeks and months on thoroughly isolating hundreds of phages is also a costly affair in regards of time and workhours. And still, also by individual isolation resulting phages may end up being similar specimen.

With the HiTS method phage libraries can be build and sequenced after just four consecutive days of sample processing. The fast turnaround is of particular importance when screening for phages for PT, but not all phages are suitable for PT. Lysogenic phages should be avoided as they do not necessarily lyse their hosts and may also increase virulence of their hosts by lysogenic conversion [43]. Some phages, also those with a lytic lifestyle, code for genes with unwanted genetic traits such as toxins, superantigens, intracellular survival/host cell attachments proteins or ARGs which can be spread to bacterial communities through transduction [44], [45]. This is especially relevant to consider when isolating phages from wastewater, as treatments plants can be considered hotspots for ARGs [45]. Fortunately, ARGs and other unwanted genetic traits can, for a large part, be deduced by genetic analyses and thus phages coding for them can with decent confidence be excluded. Hence, the HiTS method allows selection of new candidate phages after a few weeks of screening, sequencing and analysing. The ability of the candidate phages to infect the target host is already verified and the absence of undesired genetic traits confirmed, consequently the phages are now ready for experimental validation and final isolation, if required.

In conclusion, the HiTS method presented here has the potential to efficiently detect the diversity of and crudely isolate phages relevant for PT and biocontrol which are abundant in the sample matrix explored. The HiTS method is simple, fast and cost-efficient. It can prove to be a valuable, scalable method in the case of urgent needs for PT suitable phages targeting specific clinical infections. With the HiTS method establishment of sizeable discovery phage banks becomes fast and efficient. Such phage discovery banks could be lifesavers eliminating the need to spend time on isolating new PT phages and would also facilitate important phage taxonomy and ecology studies and can be explored for industrially-relevant biotechnological applications.

## Supplementary Materials

The following are available online at www.mdpi.com/xxx/s1, Figure S1: Illustration of Checker plating, Table S1: List of samples, Table S2: List of unique *P. aeruginosa* phages, Table S3: List of unique *Enterococcus* phages, Table S4: List of unique *Salmonella* phages, Figure S2: Phylogenomic nucleotide distances of the 39 unique *P. aeruginosa* phages, Figure S3: Phylogenomic nucleotide distances of the four unique *Enterococcus* phages, Figure S4: Rarefaction curves and diversity indices, Figure S6: Phylogenomic amino acid distances of the 59 unique *Salmonella* phages.

## Author Contributions

Conceptualization, N. S. O., W. K.: and L. H. H.; methodology, N. S. O., W. K. and L. H. H.; validation, N. S. O., W. K., N. B. H. and L. H. H.; formal analysis, N. S. O.; investigation, N. S. O.; resources, N. S. O. and L. H. H.; data curation, N. S. O.; writing—original draft preparation, N. S. O.; writing—review and editing, N. S. O., W. K., N. B. H. and L. H. H.; visualization, N. S. O.; supervision, N. S. O., W. K. and L. H. H.; project administration, N. S. O., W. K. and L. H. H.; funding acquisition, W. K. and L. H.H.

## Acknowledgments

This study had not been possible without the much-appreciated contribution by the many collaborating members of the Danish Water and Wastewater Association (DANVA), who kindly supplied us with time-series of wastewater samples from their treatment facilities. A special thanks to the Billund and Grindsted treatment facilities of Billund Vand & Energi, Lynetten, Avedøre and Damhusåen treatment facilities of BIOFOS, Kolding treatment facility of BlueKolding, Esbjerg Øst, Esbjerg Vest, Ribe, Varde og Skovlund treatment facilities of DIN Forsyning, Drøsbro, Hadsten, Hammel, Hinnerup and Voldum treatment facilities of Favrskov Forsyning, Haerning treatment facility of Herning Vand, Hillerød treatment facility of Hillerød Forsyning,Lemvig treatment facility of Lemvig Vand og Spildevand, Bevtoft, Gram, Haderslev, Halk, Jegerup, Nustrup, Over Jerstal, Skovby, Skrydstrup, Sommersted, Vojens and Årøsund treatment facilities of Provas, Marselisborg, Egå and Viby treatment facilities of Aarhus vand, Hedensted, Juelsminde and Tørring treatment facilities of Hedensted Spildevand, Ejby Mølle, Bogense, Hofmansgave, Hårslev, Nordvest, Nordøst, Otterup and Søndersø treatment facilities of VandCenterSyd, Øst and Vest treatment facilities of Aalborg Forsyning and finally Helsingør treatment facility of Forsyning Helsingør.

## Supplementary Figures and Tables for

**Figure S1.**
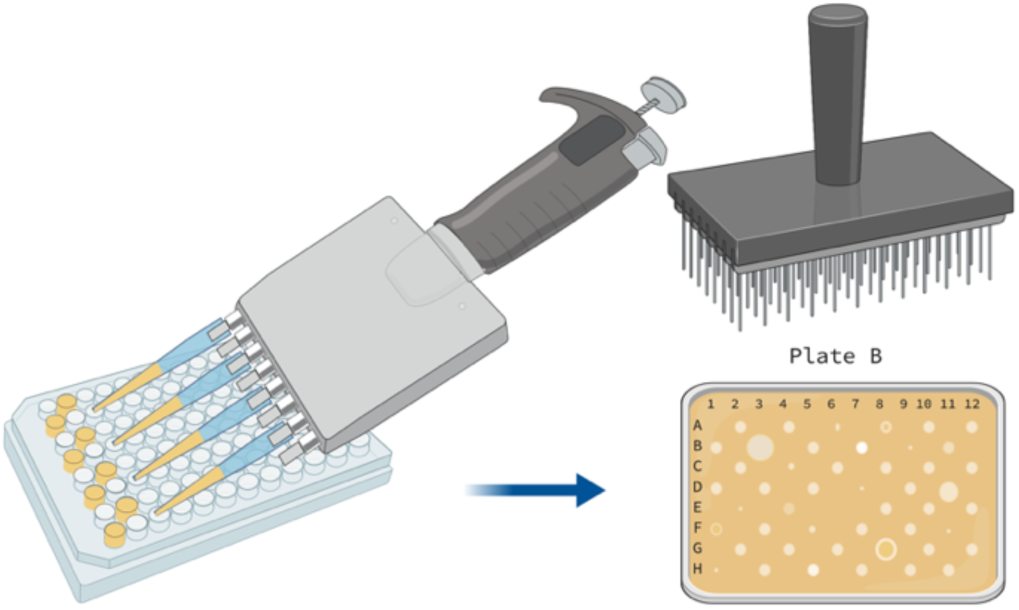
Illustration of chequer plating, removing every second row of pipette-tips enables the easy transfer of lysate to every second well of a well-plate in a chequer patter ensuring a safe distance between clearing zones when plating. Illustration created in BioRender.

**Table S1.**
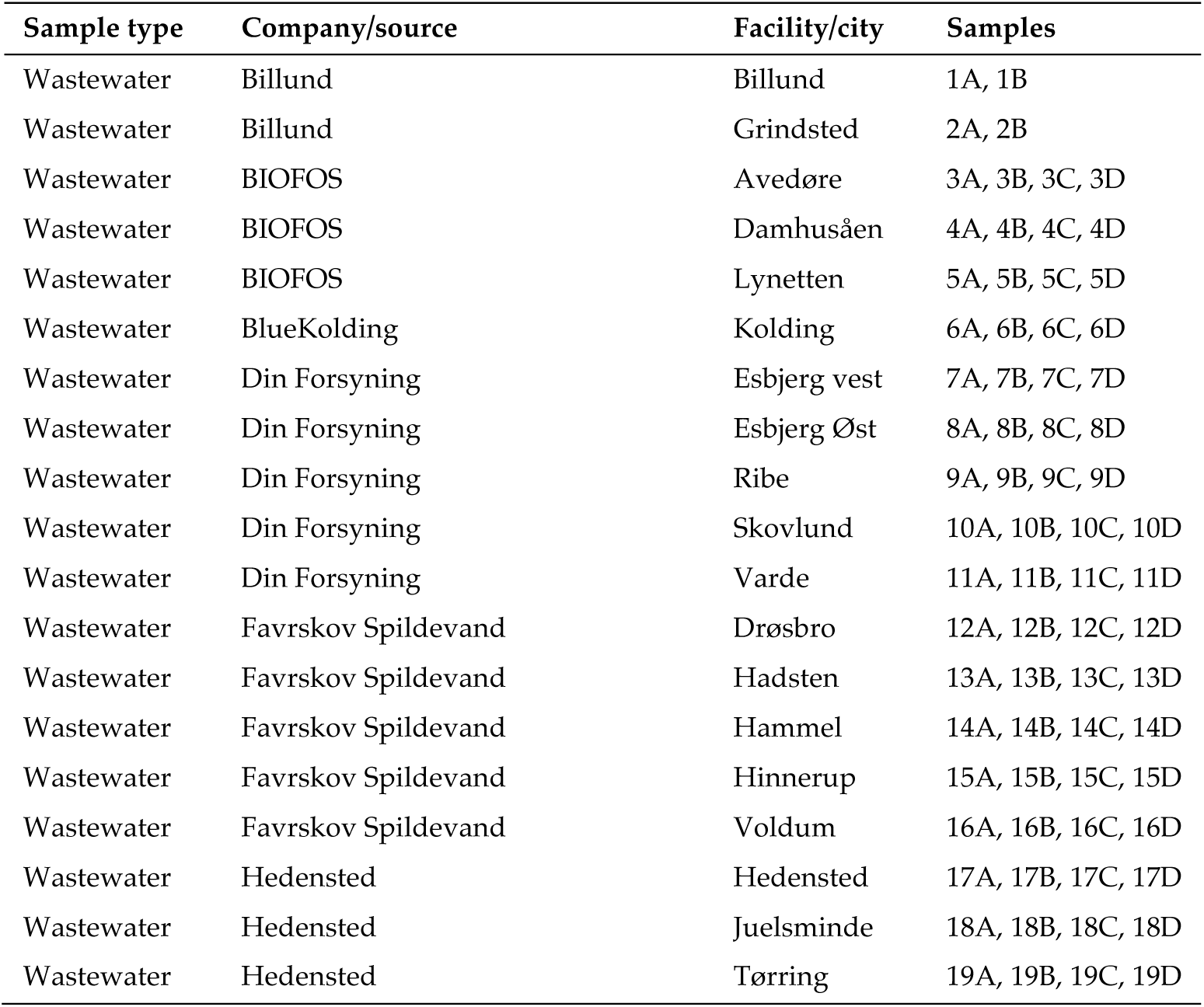
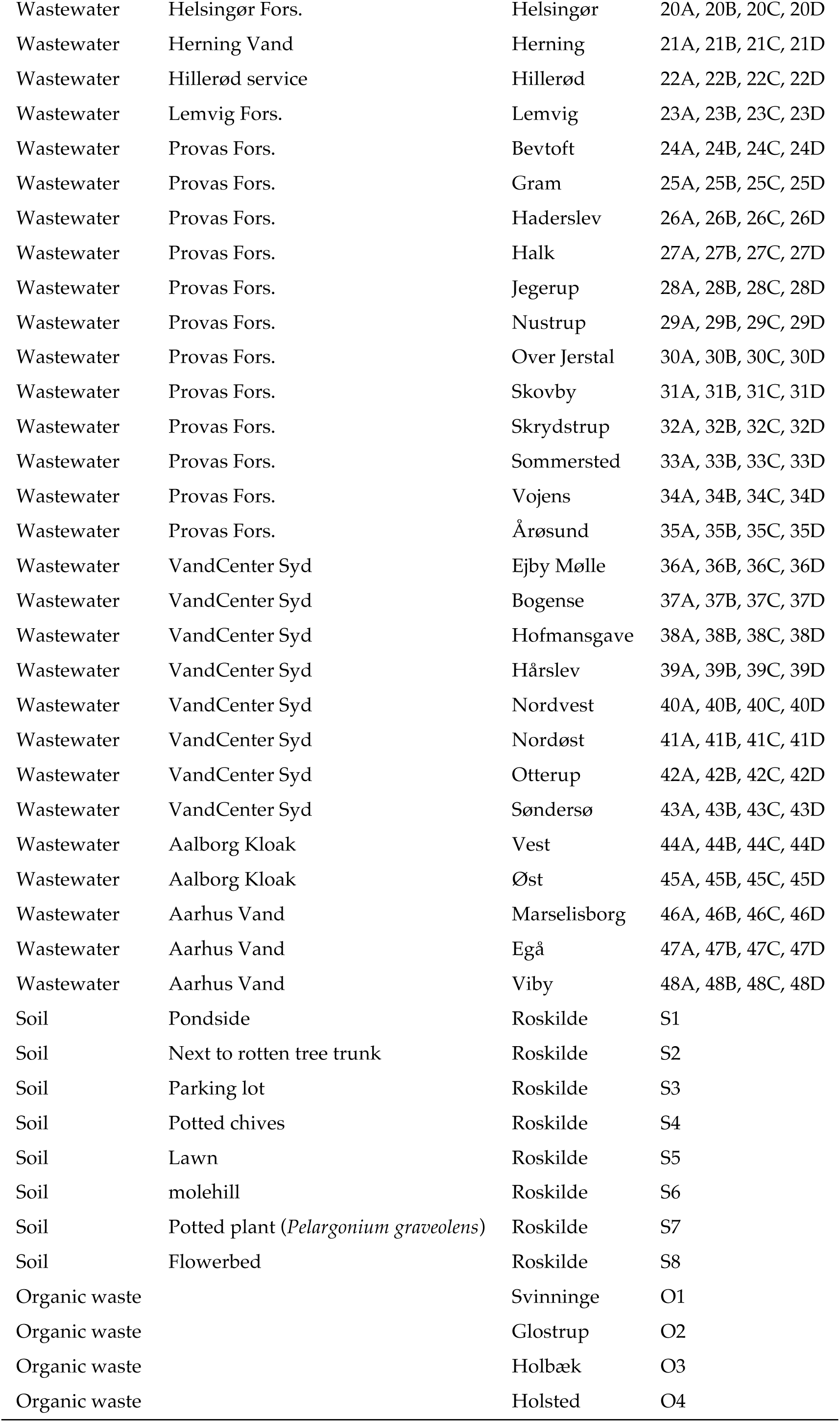
List of samples used for phage screenings.

**Table S2.** List of unique *P. aeruginosa* phages i.e. those which differ by >5% from other phages in the dataset, identified in the LV screening of Danish wastewater, soil and organic waste.

| phage | genome size<br>(bp) | GC<br>(%) | CDS<br>(n) | tRNAs<br>(n) | Novel<sup>1</sup> | taxonomy | Sample | Accession |
| --- | --- | --- | --- | --- | --- | --- | --- | --- |
| clash | 44912 | 52.1 | 68 | 3 | * | *Podoviridae*;<br>*Bruynoghevirus* | 1A, 2B, 9A | MT119362 |
| otherone | 44930 | 52.1 | 68 | 3 |  | *Podoviridae*;<br>*Bruynoghevirus* | 1B | MT119373 |
| oldone | 45313 | 52.5 | 69 | 4 | * | *Podoviridae*;<br>*Bruynoghevirus* | 4C | MT119371 |
| Iggy | 60769 | 56.5 | 90 | - |  | *Siphoviridae*. | 4D | MN029011 |
| datas | 60746 | 54.8 | 89 | - |  | *Myoviridae*; *Pbunavirus* | 21B | MT119378 |
| goonie | 64599 | 55.6 | 94 | - |  | *Myoviridae*; *Pbunavirus* | 8C | MT133561 |
| elmo | 65276 | 55.0 | 91 | - |  | *Myoviridae*; *Pbunavirus* | 4B | MT119364 |
| willy | 65355 | 55.6 | 92 | - |  | *Myoviridae*; *Pbunavirus* | 20C | MT133562 |
| billy | 65580 | 54.9 | 91 | - |  | *Myoviridae*; *Pbunavirus* | 6D | MT133563 |
| steven | 65632 | 54.9 | 92 | - |  | *Myoviridae*; *Pbunavirus* | 11D | MT119370 |
| antinowhere | 65852 | 55.2 | 91 | - | * | *Myoviridae*; *Pbunavirus* | 2B, 5D | MT119374 |
| misfit | 65887 | 54.9 | 93 | - |  | *Myoviridae*; *Pbunavirus* | 3B | MT119367 |
| chumba | 65852 | 55.0 | 92 | - |  | *Myoviridae*; *Pbunavirus* | 3D, 7D, 8D | MT119375 |
| goodold | 66226 | 55.6 | 95 | - |  | *Myoviridae*; *Pbunavirus* | 6A, 6B, 9A | MT119365 |
| chunk | 66231 | 55.6 | 93 | - |  | *Myoviridae*; *Pbunavirus* | 1B | MT119376 |
| debbie | 66259 | 55.7 | 93 | - |  | *Myoviridae*; *Pbunavirus* | 7A, 8B, 10A, 12D,<br>15C, 18C, 18D,<br>21C, O1, O2, O3 | MT119363 |
| jett | 66262 | 55.6 | 93 | - | * | *Myoviridae*; *Pbunavirus* | 8A, 11A, 13D | MT119366 |
| crassa | 66295 | 55.2 | 91 | - | * | *Myoviridae*; *Pbunavirus* | 4A | MT119377 |
| cory | 66395 | 55.2 | 91 | - | * | *Myoviridae*; *Pbunavirus* | 4C | MT119372 |
| sortsol | 66445 | 55.7 | 91 | - | * | *Myoviridae*; *Pbunavirus* | 4D, 13B, 19A | MT119369 |
| shane | 66467 | 55.6 | 95 | - |  | *Myoviridae*; *Pbunavirus* | 14D, 16B | MT119368 |
| fnug | 278899 | 36.9 | 360 | 6 | * | *Myoviridae*; *Phikzvirus* | 1B | MT133560 |

<sup>1</sup>Novel phage species classified by a 95% nucleotide similarity demarcation are indicated by an asterisk.

**Table S3.** List of unique *Enterococcus* phages i.e. those which differ by >5% from other phages in the dataset, identified in the SV screening of Danish wastewater

| Phage | genome size<br>(bp) | GC<br>(%) | CDS<br>(n) | tRNAs<br>(n) | Novel<sup>1</sup> | Taxonomy | Sample | Accession |
| --- | --- | --- | --- | --- | --- | --- | --- | --- |
| heks | 39708 | 35.0 | 64 | - | * | *Siphoviridae*; *Efqatrovirus* | 4A | MT119359 |
| Nonaheksakonda | 41994 | 34.6 | 74 | - | * | *Siphoviridae*; *Efqatrovirus* | 45D | MK125140 |
| vipetofem | 85371 | 30.3 | 134 | 1 |  | *Siphoviridae* | 44D | MT119361 |
| nattely | 85669 | 30.2 | 131 | 1 | * | *Siphoviridae* | 46D | MT119360 |

<sup>1</sup>Novel phage species classified by a 95% nucleotide similarity demarcation are indicated by an asterisk.

**Table S4.**
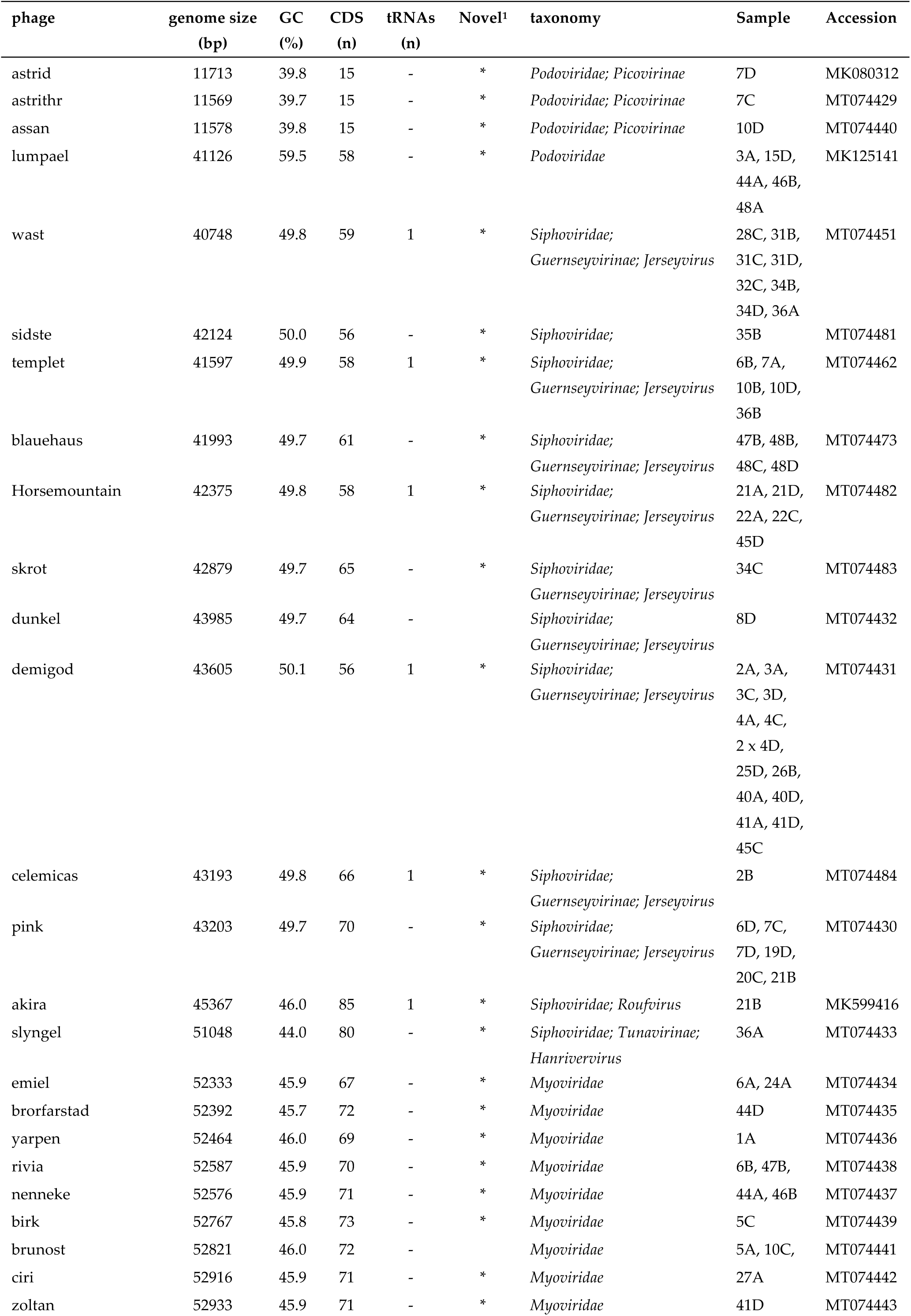
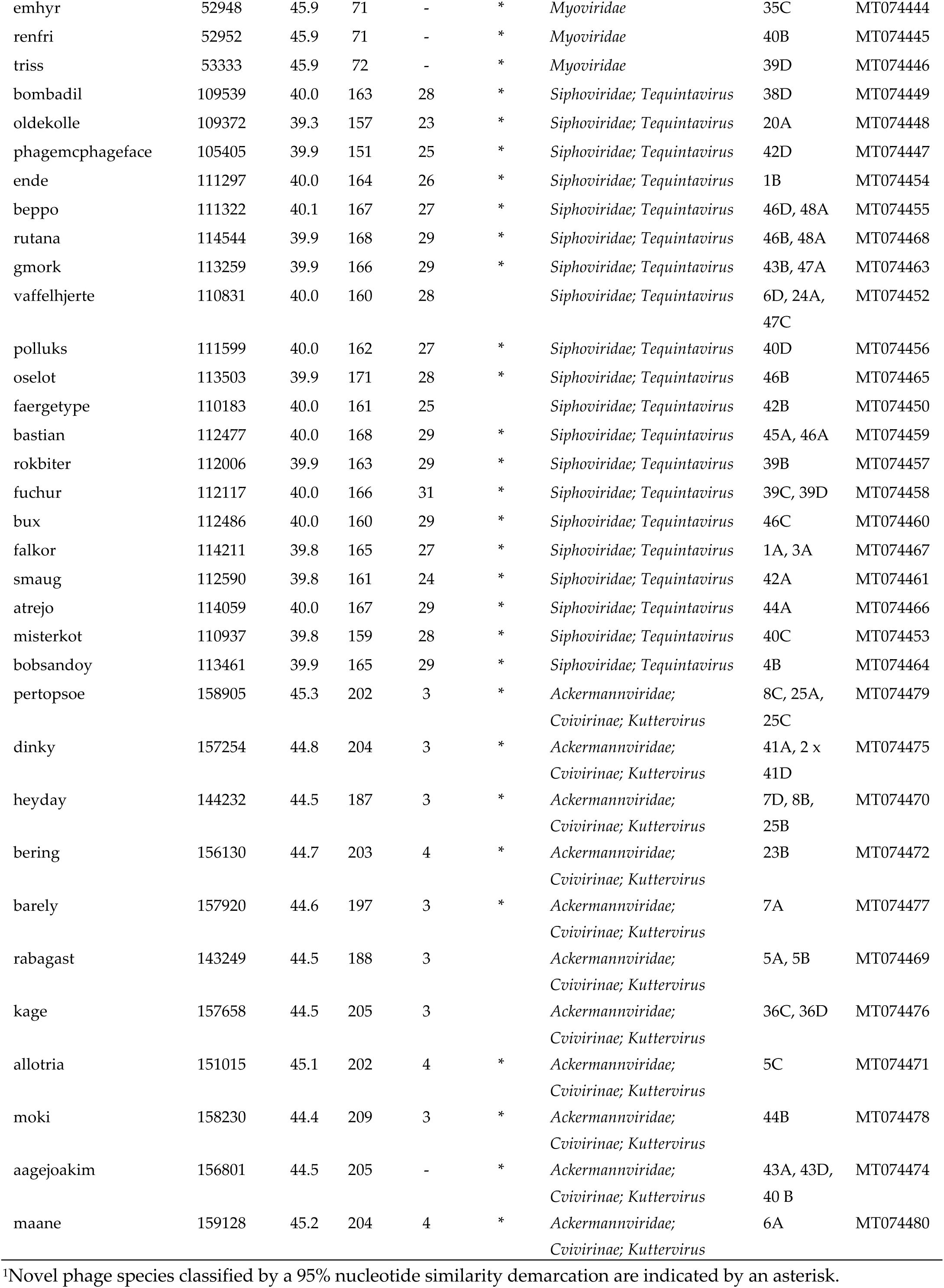
List of unique *Salmonella* phages i.e. those which differ by >5% from other phages in the dataset, identified in the SV and LV screenings of Danish wastewater.

**Figure S2.**
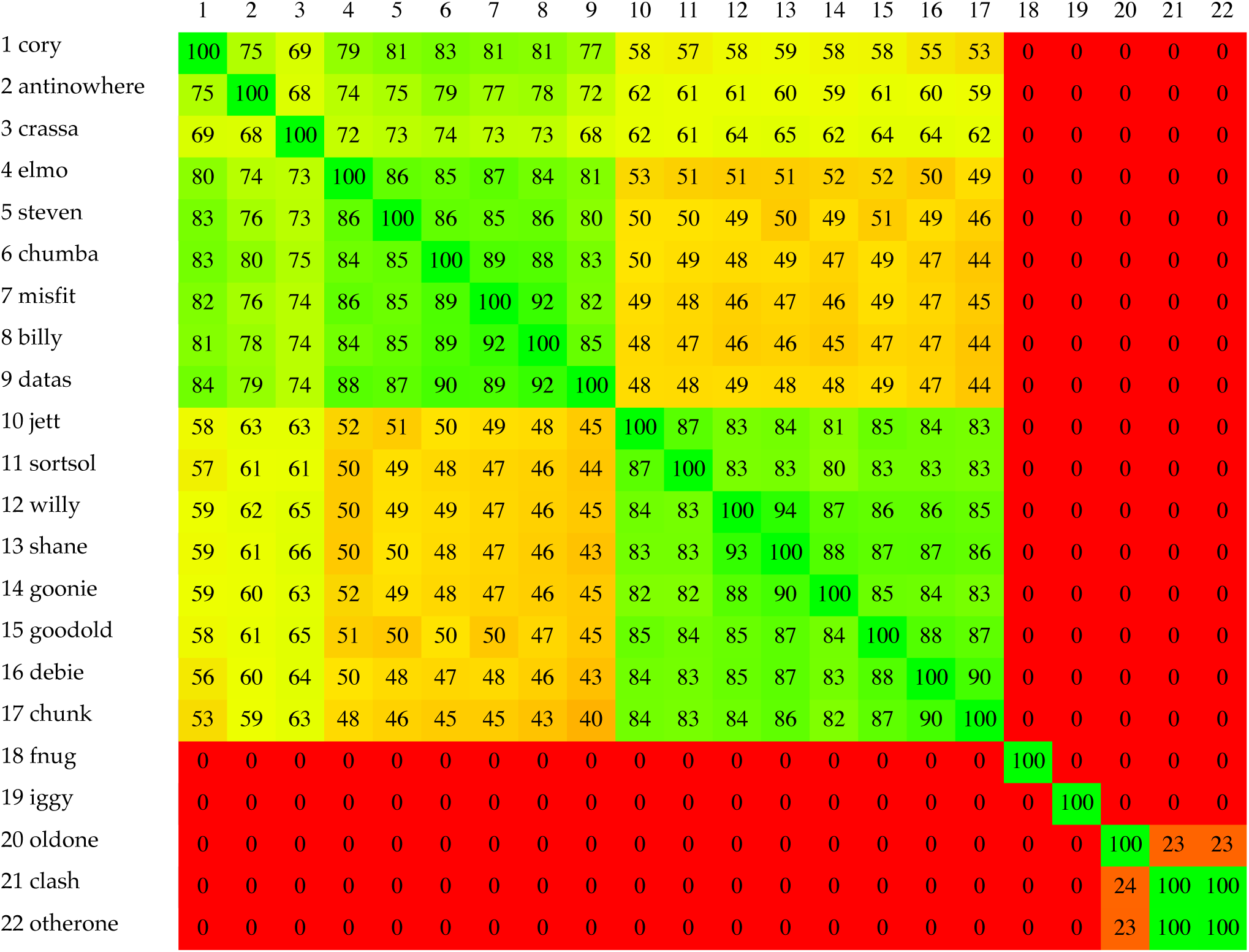
phylogenomic nucleotide distances of the 39 unique *P. aeruginosa* phages (Gegenees, BLASTn: fragment size: 200, step size: 100, threshold: 0).

**Figure S3.**
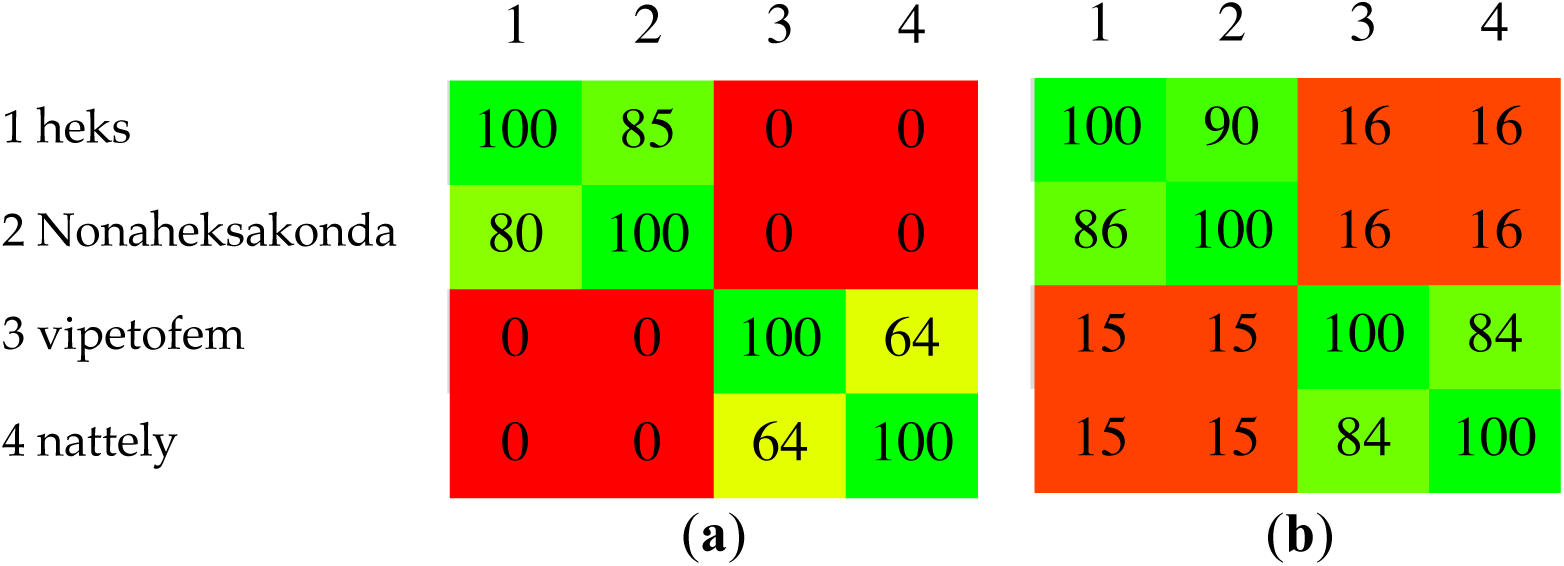
(**a**) Phylogenomic nucleotide distances of the 4 unique *Enterococcus* phages phages (Gegenees, BLASTn: fragment size: 200, step size: 100, threshold: 0%). (**b**) Phylogenomic amino acid distances of the 4 unique *Enterococcus* phages (Gegenees, BLASTx: fragment size: 200, step size: 100, threshold: 0%).

**Figure S4.**
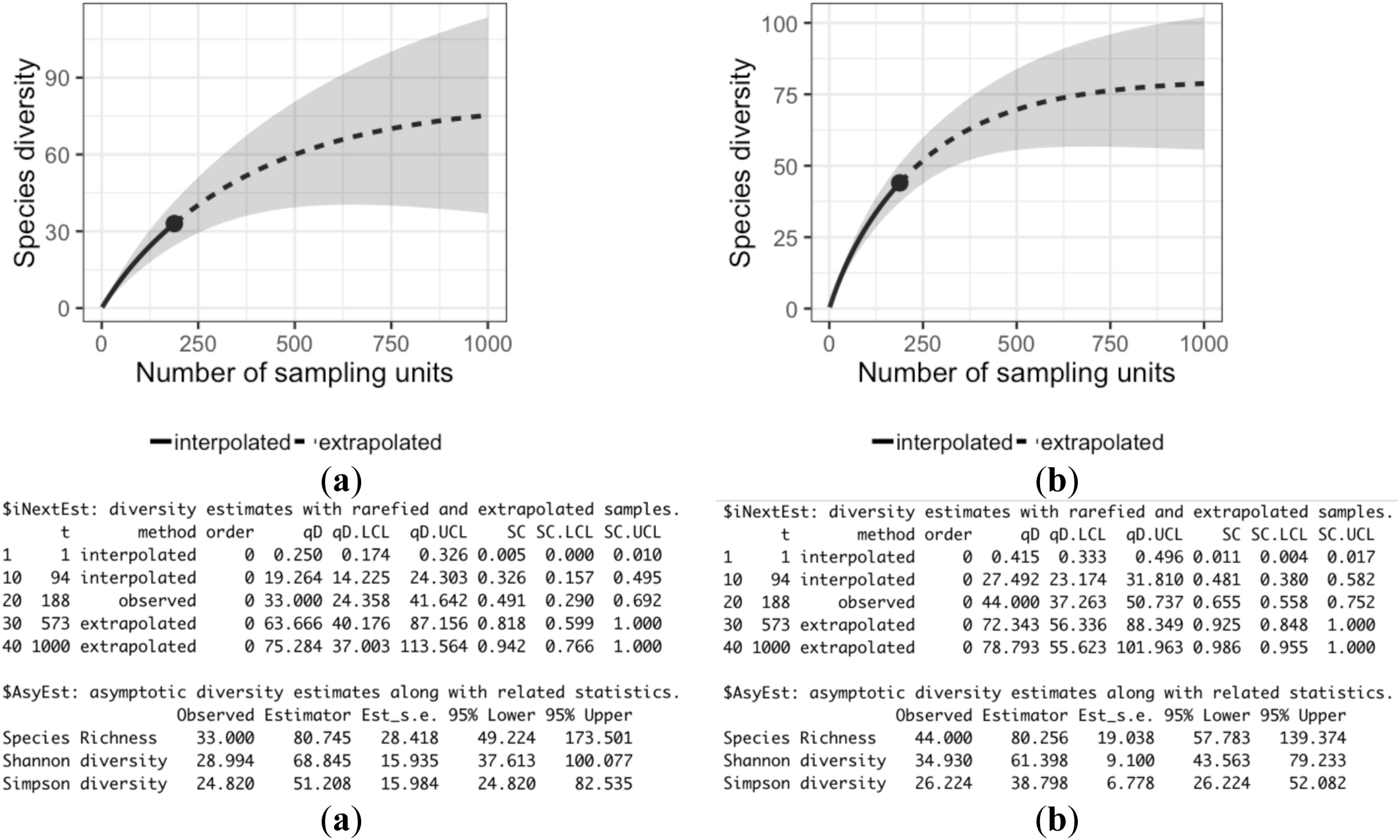
Rarefaction curves for the *S. enterica* screenings; (**a**) small volume (SV) screening and (**b**) large volume (LV) screening. (**c**) Diversity indices for the small volume (0.5ml) screening based on identification of phages species in all 188 wastewater samples. (**d**) Diversity indices for the large volume (0.5ml) screening based on identification of phages species in all 188 wastewater samples.

**Figure S5.**
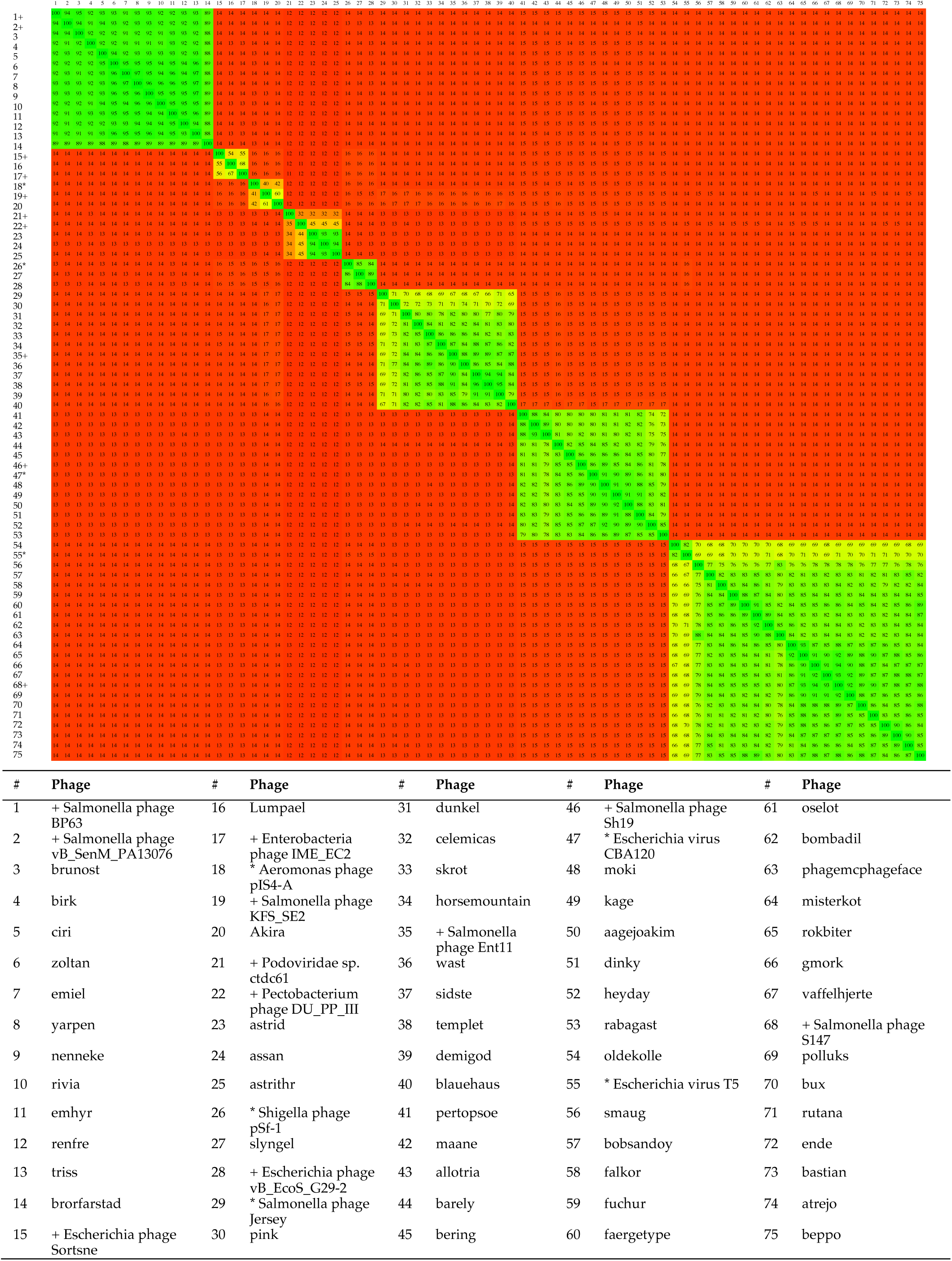
Phylogenomic amino acid distances of the 59 unique *Salmonella* phages, selected closest relatives and type species of respective genera (Gegenees, BLASTx: fragment size: 200, step size: 100, threshold: 0%). Type species are denoted by an asterisk (*), close relative by plus (+).

